# On the stock structure bias of the space-time fidelity of mark-recapture studies

**DOI:** 10.64898/2026.05.14.725068

**Authors:** Lars Witting

## Abstract

Mark-recapture analyses on the delineation of natural populations between areas often assume random sampling, with a between/within (B/W) area resighting ratio that declines towards zero as the population components of two areas become more-and-more isolated from one another, with fewer-and-fewer individuals mixing between areas. I use an individual based population model split in two areas to simulate this result, analysing also for the potential effects of the space-time fidelity of the mark-recapture sampling in the areas. I find that small B/W resighting ratios—that traditionally is taken as evidence of population isolation—can easily be observed within a completely mixing population if a random sampling scheme is restricted in space and/or time. Random sampling within restricted areas and time windows is not sufficient to estimate mixing rates and population isolation between areas, unless the resighting rates are analysed by a method that accounts both for the space-time fidelity of the scientific sampling scheme and the space-time fidelity of the distributional behaviour of the individuals in the population.

## 1 Introduction

Mark-recapture studies of individual animals, including photo resighting studies of humpback whales, are often used to delineate natural populations. This is usually done by comparing the recaptures/resightings within and across two areas assuming random sampling within each area. If individual animals move more-or-less at random within and across the two areas, the limit ratio of the proportion of resighted individuals within and across the two areas approach unity. But as the mixing of individuals between the two areas decline the number of within-area resightings start to outnumber the number of between-area resightings (Fig. 1, top plot). The ratio of the two resighting rates (bottom plot) is thus apparently an informative measure of the degree of population isolation between areas.

**Figure 1.**
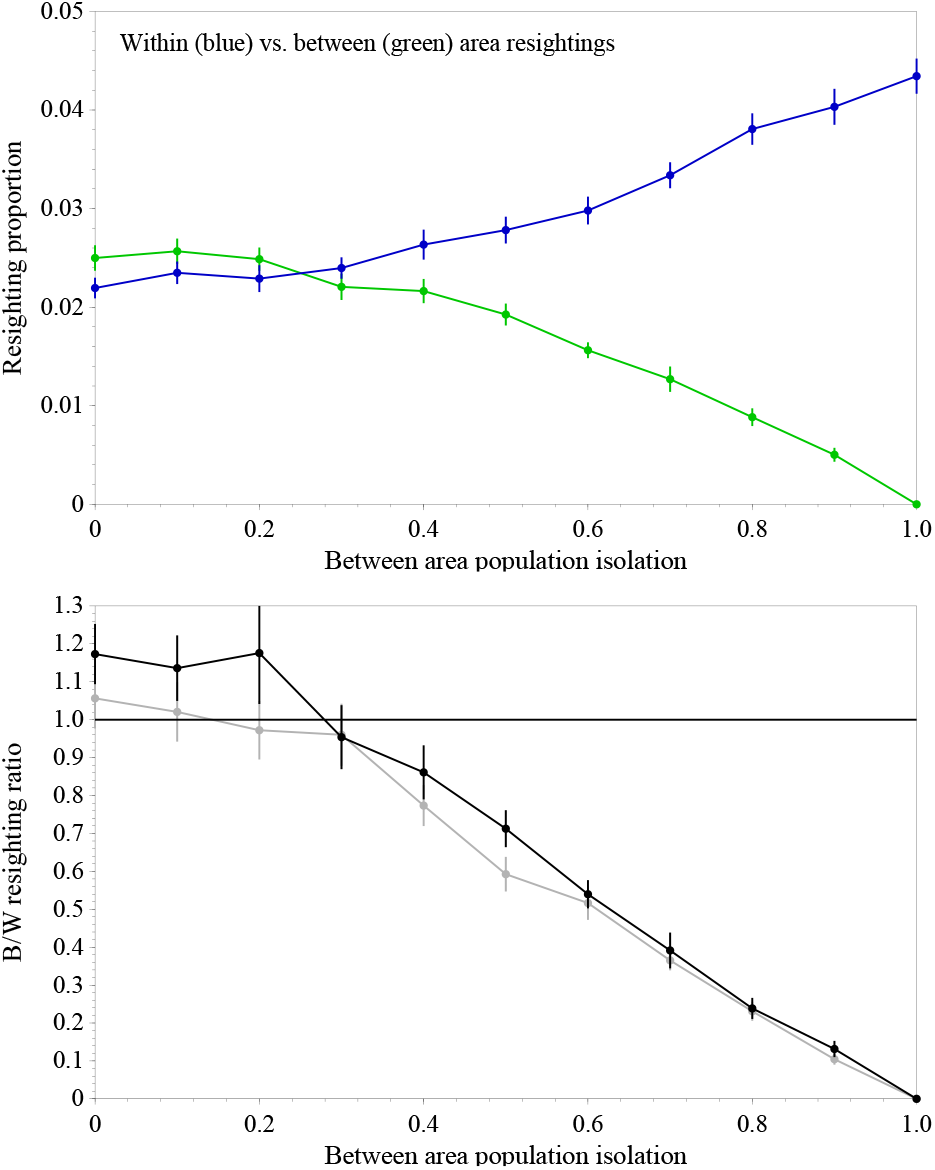
**Top Plot:** The within (blue) vs. between (green) area randomly resighted individuals (relative to total nr. of sighted individuals) as a function of the isolation between two simulated population areas with 5,000 individuals in each. **Bottom plot:** The between/within (B/W) area resighting ratio (green curve divided by blue) given individuals with no (black curve & top plot) or complete (grey) space-time fidelity (.see methods/results). A completely mixing population has no isolation boundary (*b* = 0) having individuals that mix with probability *p*_*m*_ = (1 − *b*)*/*2 = 0.5 into the non-resident area. Completely isolated populations have hard isolation boundaries (*b* = 1) and individuals with zero mixing probabilities (*p*_*m*_ = 0). The limit value of the B/W resighting ratio is one for completely mixing populations, but slightly larger in the simulation where within-year resightings are excluded only within areas.

The underlying assumption of random sampling may not be explicitly addressed in population structure studies, reflecting perhaps a deeper implicit assumption of individuals that shuffle their distributional behaviour between years. So why should we not believe the results if the within-year resightings within each area are excluded from the set of resightings.

With the present study I build an individual-based two area population model to examine the effects that the space-time fidelity of sampling has on the population structure results of mark-recapture studies. Just as the individuals of the species we study may have site and time fidelity preferences for being in particular areas at specific times of the year, so are mark-recapture population structure studies rarely sampling the complete area of the population evenly year-round. Sampling may instead have a point-like spatial distribution, as for many photo-identification photos of hump-back whales (Fig. 2) often taken from whale watching tours from a limited number of towns. Dedicated sampling by biologists may occur over larger areas, but they may still cover only a fraction of the distribution area and be logistically repeated in the same areas in the same month each year. Either way, by analysing under the assumption of random sampling we need some geographically or behaviourally imposed limited mixing of individuals between the two areas to explain the frequent observation where within-area resightings are much more common than between-area resightings (e.g. Stevick et al. 2006; Lubansky et al. 2026). As illustrated for this case in Fig. 1, site fidelity in the spatial movements of individuals is not—in the absence of a mixing boundary—sufficient to explain the dominance of within-area resightings relative to between-area resightings. But may the dominance of within-area resightings reflect the space-time fidelity of sampling?

**Figure 2.**
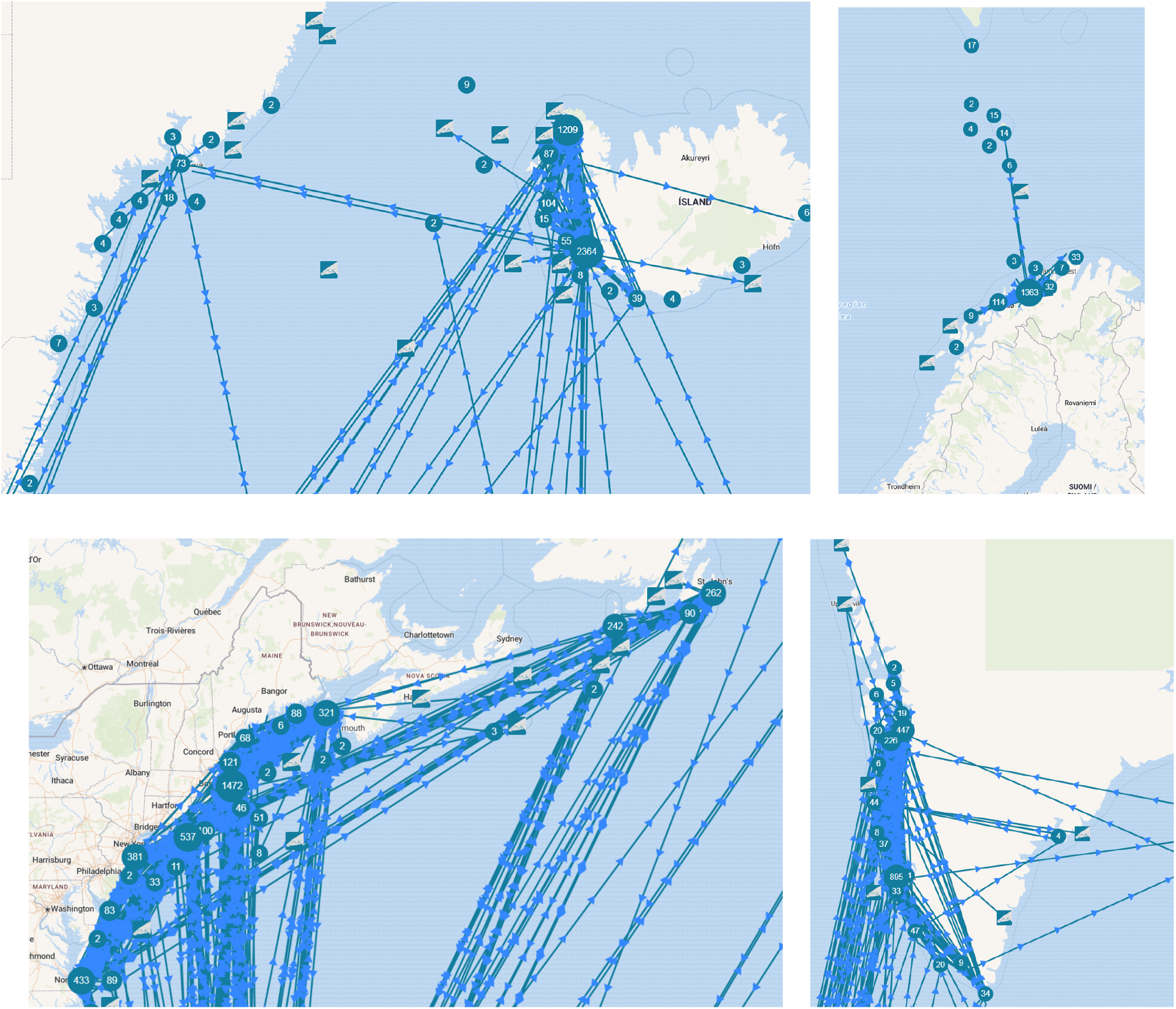
Plots from happywhale.com of positions of photos taken of humpback whales on summer feeding grounds off East Greenland and Iceland (top left), Norway (top right), Eastern North America (bottom left), and West Greenland (bottom right), with numbers showing the number of encounters from 2010 (or 2020 America) to the present. There are two major point-like sampling sites off Iceland and West Greenland, one off Norway and East Greenland, and in the order of seven off Eastern North America.

## 2 Method

To study the effect of sampling fidelity, I divide an individual-based population model with a fixed number of *n* individuals with no mortality into two components (*n*_*a*_ *& n*_*b*_) that are pre-oriented towards two (*a*_*a*_ *& a*_*b*_) non-overlapping distribution areas (*a*). Dependent upon the degree of mixing, area *a*_*a*_ individuals may be pre-disposed to being just as much in area *a*_*b*_ as in area *a*_*a*_, or pre-disposed to being mainly or only in area *a*_*a*_, but never pre-disposed to being mainly or only in area *a*_*b*_ (the same applies with opposite signs to area *a*_*b*_ individuals).

The spatial distribution behaviour of each individual is obtained by drawing behaviour randomly from the pre-disposed possibilities of behaviours as defined by the parameters of my model. The pre-disposed behaviour is identical for all individuals assigned to the same area, with the *a*_*a*_ and *a*_*b*_ area individuals having symmetrical pre-disposed behaviour in my model example that relates to 10, 000 humpback whales on half-year summer feeding grounds in the North Atlantic (*n*_*a*_ = *n*_*b*_ = 5, 000). A central component of the individual behaviour is the probability of mixing into the non-resident area as it defines the mixing rate of individuals between areas.

### 2.1 Mixing probability

To simulate the mixing between areas I model the yearly space-time distribution of each individual by a time-vector of length *n*_*t*_ that specifies where the individual is during the summer feeding period of the year, defined by 26 consecutive weeks in my example. The yearly space-time distribution is re-considered by each individual each year, with the area visited in a given week being either the same by fidelity across all years or a new area drawn at random from the pre-disposed behaviour, dependent upon the fidelity behavioural of the individual. The fidelity weeks and their fixed distributional behaviour are drawn from the pre-disposed behaviour, but only once during the model construction of each individual and kept constant thereafter.

For the initial choice of fidelity weeks, and yearly choice of non-fidelity weeks, the first choice that each individual makes is whether to go to area *a*_*a*_ or *a*_*b*_, as defined by the mixing probability

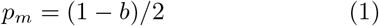

that an area *a*_*a*_ individual goes to area *a*_*b*_, or an area *a*_*b*_ individual goes to area *a*_*a*_, with *b* being the boundary parameter of isolation between the two population components, with *b* = 0 being no boundary with full mixing (*p*_*m*_ = 0.5) and no population isolation, and *b* = 1 a hard boundary with complete isolation and no mixing (*p*_*m*_ = 0) of individuals between the two areas.

### 2.2 Individual fidelity

The next choice of the individual is to decide where to go in the chosen area *a*_*a*_ or *a*_*b*_, and for this I partition each of the two areas into a pre-defined number (*n*_*a,s*_) of sub-areas *a*_*a,s*_ and *a*_*b,s*_. For my example, I have 260 sub-areas for each main area, implying that an individual can potentially cover ten percent of a given area, or five percent of the combined area, in half a year of feeding. The choice of sub-area visited in each week is again drawn at random from a pre-disposed behaviour that allows the individual to potentially choose among all the sub-areas in the main area *a*_*a*_ or *a*_*b*_.

The way sub-areas are chosen is slightly different for the initial choice of the fidelity weeks, and the yearly choices of the non-fidelity weeks. For this I define the pre-disposed fidelity behaviour by a fidelity parameter 0 ≤ *f* ≤ 1, where *f* = 0 defines random behaviour and *f* = 1 complete space-time fidelity, i.e., individuals that always go to the same areas in the same weeks each year. This parameter defines

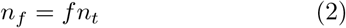

fidelity weeks per year, and these are positioned at random in the space-time vector of each individual, with the constantly visited sub-area of each fidelity week being decided from a random mixing decision followed by a random sub-area decision among all the sub-areas of the chosen main area, allowing the sequence of independent draws to potentially draw the same sub-area for all fidelity weeks.

The fidelity parameter is not only determining the number of fidelity weeks with fixed sub-areas to visit, but it is also making the individual more behaviourally restricted in the potential sub-areas that are considered for the remaining *n*_*t*_ − *n*_*f*_ weeks where the individual decides each year where to go. This is done by defining a vector of randomly drawn potential sub-areas from which the visited areas are drawn at random, with the length of the potential sub-area vector being

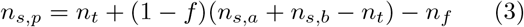

with the sub-areas of the vector being determined by random mixing draws followed by random sub-areas drawn from the complete vector of sub-areas of the mixing drawn main area. This vector of potential sub-areas to visit at random contracts to zero length for completely space-time fidelity behaving individuals (*f* = 1 with *n*_*f*_ = *n*_*t*_ *& n*_*s,p*_ = 0), and expands to potentially include all sub-areas for random behaviour with no space-time fidelity (*f* = 0 with *n*_*f*_ = 0 and *n*_*s,p*_ = *n*_*s,a*_ + *ns, b*).

### 2.3 Space-time fidelity of sampling

Having a model population where 10, 000 individuals potentially repositions themselves among 2 × 260 sub-areas each week over a 26-week summer feeding season each year, I mark/photo identify/sample *n*_*a,id*_ unique individuals per year from each main area *a*_*a*_ and *a*_*b*_ for a number of years. This sampling is structured by the space-time fidelity of the sampling scheme that determines where among the 2 × 260 sub-areas, and when across the 26 weeks, the sampling occurs, with random sampling within each of the chosen sub-areas during the chosen weeks continuing until *n*_*a,id*_ unique individuals are identified per year.

For this I define a time-fidelity (*f*_*a,t*_) and site-fidelity (*f*_*a,s*_) parameter for the sampling in each main area, having identical sampling schemes in the two areas in my example. The time-fidelity

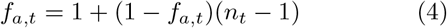

defines the duration of the sampling season from one to *n*_*t*_ = 26 weeks, and the site-fidelity

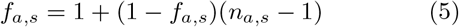

defines the spatial extend of the sampling ranging from one to the *n*_*a,s*_ = 260 sub-areas of each main area. The actual weeks and sub-areas of the sampling are assigned randomly and maintained across all sampling years, with the yearly sampling of *n*_*a,id*_ unique individuals occurring randomly across the chosen sub-areas and weeks of the sampling scheme. In my example I sample 50 unique individuals from each main area each year covering 10 years of sampling, and run a number of independent iterations for each parametrisation to estimate the average and standard error of the resighting numbers.

## 3 Results

The case with random sampling in space and time is shown in Fig. 1, with the top plot showing the proportion of the within-area (blue curve) and between-area (green curve) resighted individuals (relative to the total number of unique individuals sighted within and between areas) as a function of the mixing boundary of population isolation, given otherwise randomly behaving individuals with no space-time fidelity. The green curve is divided by the blue curve to provide the between/within (B/W) area resighting ratios in the bottom plot, with randomly behaving individuals (*f* = 0) represented by the black curve, and complete space-time fidelity behaviour (*f* = 1) represented by the grey curve (not shown in the top plot). Both curves are similar showing an approximately linear drop in the B/W resighting ratio from about one for a complete mixing population (*b* = 0) to zero for completely isolated populations with hard population boundaries (*b* = 1) and no mixing. This follows the expectation that we should be able to successfully estimate the mixing between populations given random sampling from both populations; a result that holds independently of the presence versus absence of space-time fidelity in the behaviour of the studied individuals.

Fig. 3 expands the case of a completely mixing population (left-most limit of Fig. 1) as a function of the space-time fidelity of the individuals in the population, with the black dots approaching the expected invariant B/W resighting ratio around unity when sampling is random. The B/W resighting ratio however drops considerably when sampling is non-random, becoming a declining function of the space-time fidelity of the individuals in the population. The red curve, that represents a point-like random sampling within a single sub-area in each main area over the whole season, has a B/W resighting ratio that starts around 60% for randomly behaving individuals and drops below 5% for individuals with strong space-time fidelity. The blue curve illustrates that sampling with a 5% area coverage over four weeks has a slightly higher B/W resighting ratio than point-like sampling over the whole season (dropping to 10%), while the green curve with 25% area coverage over four weeks maintains a B/W ratio above 35% for individuals with strong space-time fidelity. Similar relationships, but with a stronger dominance of within-areas resightings, are shown in Fig. 4 for cases with 50% and 75% population isolation, corresponding to mixing probabilities/rates of 25% and 12.5%.

**Figure 3.**
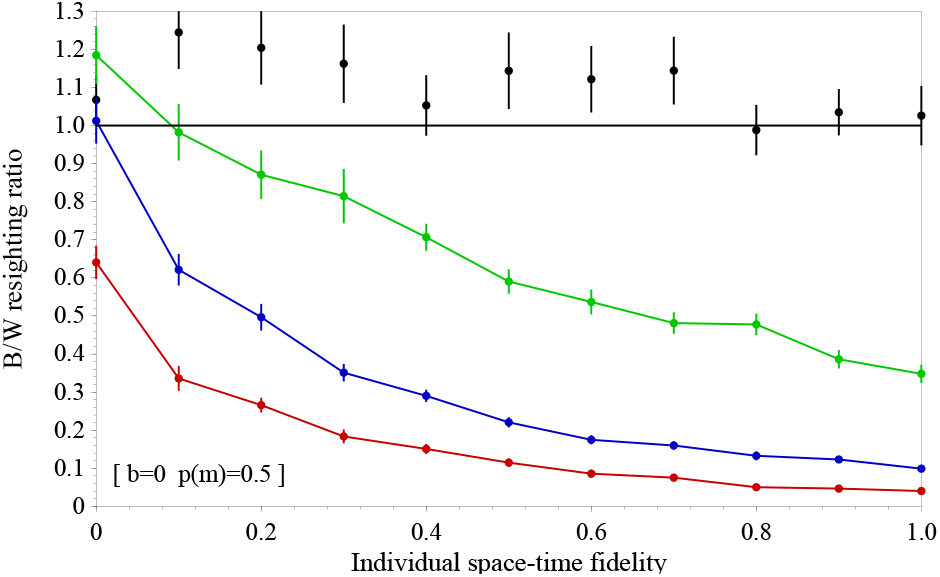
The B/W area resighting ratio as a function of the space-time fidelity of individuals within a single population with complete mixing (*p*_*m*_ = 0.5) and no internal isolation boundaries (*b* = 0). **Black:** Random sampling across the complete distribution area during the complete half-year season. **Green:** Random sampling across 25% of the area covering one month. **Blue:** Random sampling across 5% of the area covering one month. **Red:** Local sampling during the complete half-year season.

**Figure 4.**
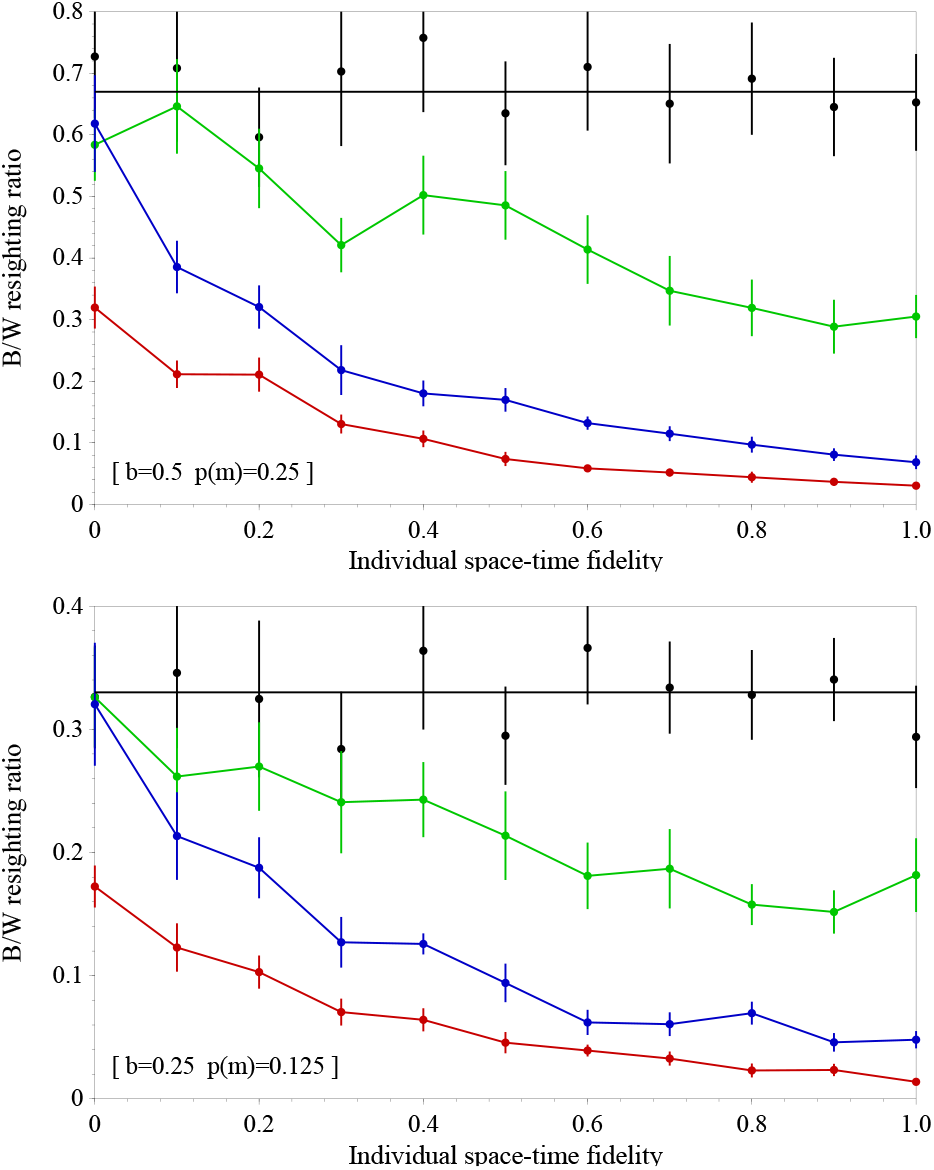
The B/W area resighting ratio as a function of the space-time fidelity of individuals for 50% (top plot) and 75% (bottom) population isolation, corresponding to individual mixing probability of 0.25 and 0.125. **Black:** Random sampling across the complete distribution area and half-year season. **Green:** Random sampling across 25% of the area covering one month. **Blue:** Random sampling across 5% of the area covering one month. **Red:** Local sampling during the complete half-year season.

## 4 Discussion

On the positive side we find that B/W resighting ratios above say 10% can only occur if individuals have at least about a five percent chance of not being in their resident area. On the negative side, B/W resighting ratios below 10% can easily be observed within a completely mixing population if a random sampling scheme is restricted in space and/or time. Random sampling within restricted areas and time windows is therefore not sufficient to estimate mixing rates between areas, unless the resighting rates are analysed by a method that accounts for the space-time fidelity of both the scientific sampling scheme and the distributional behaviour of the individuals in the population. Non-random sampling within restricted areas and time windows will only make things worse, and maybe impossible to disentangle into realistic estimates of mixing.

Facing these difficulties, it is probably in many cases better to step back and reconsider the whole concept of population components that belong to specific areas with a restricted mixing of individuals between them, unless the delineation of the areas represents a real physical migration/mixing barrier.

The delineation of areas in many population structure studies is often rather arbitrary from the point of view of the species being studied, although they often make sense from an anthropogenetic management point of view. But the structure of management is (usually) not defining the biological structure of the species, while the biological structure of the species should ideally inform management. We may therefore keep delineated areas as management units but discard them as biological entities.

## Notes

### Competing Interest Statement

The authors have declared no competing interest.

